# Genome sequence of the euryhaline Javafish medaka, *Oryzias javanicus*: a small aquarium fish model for studies on adaptation to salinity

**DOI:** 10.1101/593673

**Authors:** Yusuke Takehana, Margot Zahm, Cédric Cabau, Christophe Klopp, Céline Roques, Olivier Bouchez, Cécile Donnadieu, Celia Barrachina, Laurent Journot, Mari Kawaguchi, Shigeki Yasumasu, Satoshi Ansai, Kiyoshi Naruse, Koji Inoue, Chuya Shinzato, Manfred Schartl, Yann Guiguen, Amaury Herpin

## Abstract

**Background:** The genus *Oryzias* is constituted of 35 medaka-fish species each exhibiting various ecological, morphological and physiological peculiarities and adaptations. Beyond of being a comprehensive phylogenetic group for studying intra-genus evolution of several traits like sex determination, behaviour, morphology or adaptation through comparative genomic approaches, all medaka species share many advantages of experimental model organisms including small size and short generation time, transparent embryos and genome editing tools for reverse and forward genetic studies. The Java medaka, *Oryzias javanicus*, is one of the two species of medaka perfectly adapted for living in brackish/sea-waters. Being an important component of the mangrove ecosystem, *O. javanicus* is also used as a valuable marine test-fish for ecotoxicology studies. Here, we sequenced and assembled the whole genome of *O. javanicus*, and anticipate this resource will be catalytic for a wide range of comparative genomic, phylogenetic and functional studies.

**Findings:** Complementary sequencing approaches including long-read technology and data integration with a genetic map allowed the final assembly of 908 Mbp of the *O. javanicus* genome. Further analyses estimate that the *O. javanicus* genome contains 33% of repeat sequences and has a heterozygosity of 0.96%. The achieved draft assembly contains 525 scaffolds with a total length of 809.7 Mbp, a N50 of 6.3 Mbp and a L50 of 37 scaffolds. We identified 21454 expressed transcripts for a total transcriptome size of 57, 146, 583 bps.

**Conclusions:** We provide here a high-quality draft genome assembly of the euryhaline Javafish medaka, and give emphasis on the evolutionary adaptation to salinity.

## DATA DESCRIPTION

### Introduction/Background information

Medaka fishes belong to the genus *Oryzias* and are an emerging model system for studying the molecular basis of vertebrate evolution. This genus contains approximately 35 species, individually exhibiting numerous morphological, ecological and physiological differences and specificities (1–4). In addition, they all share many advantages of experimental model organisms, such as their small size, easy breeding, short generation time, transparent embryos, transgenic technology and genome-editing tools, with the “flag ship” species of this genus, the Japanese rice fish, *Oryzias latipes* (5, 6). Such phenotypic variations together with cutting edge molecular genetic tools make possible to identify major loci that contribute to evolutionary differences, and to dissect the roles of individual genes and regulatory elements by functional tests. For example, a recent genetic mapping approach using interspecific hybrids identified the major chromosome regions that underlie the different hyperosmotic tolerance between species of the *Oryzias* genus (7). Medaka fishes are also excellent models to study evolution of sex chromosomes and sex-determining loci among species (8–11), with the advantage of being also suitable models for providing functional evidences for these novel sex-determining genes by gain-of-function and/or loss-of-function experiments (12, 13).

Among these species, the Java medaka, *Oryzias javanicus* (Figure 1), is unique as being the prototypic species of this genus with respect to adaptation to seawater. Previous phylogenetic studies divided the genus *Oryzias* into three monopyletic groups: *(i) javanicus, (ii) latipes* and *(iii) celebensis* species groups (4, 14). Most of the *Oryzias* species inhabit mainly freshwater biotopes while only two species, which belong to the *javanicus* group, live in sea- or brackish waters. One is *O. javanicus*, found in mangrove swamps from Thailand to Indonesia, and the other is *O. dancena* (previously named *O. melastigma*) living both in sea- and freshwaters from India to Malaysia. Although both species are highly adaptable to seawater, *O. javanicus* prefers hyperosmotic conditions while *O. dancena* favours hypoosmotic conditions at the west coast of Malaysian peninsula where their distribution ranges overlap (15). In addition, *O. javanicus* is an important component of the mangrove ecosystem (16), and has been used as a valuable marine test fish in several ecotoxicology studies (17, 18).

**Figure 1:**
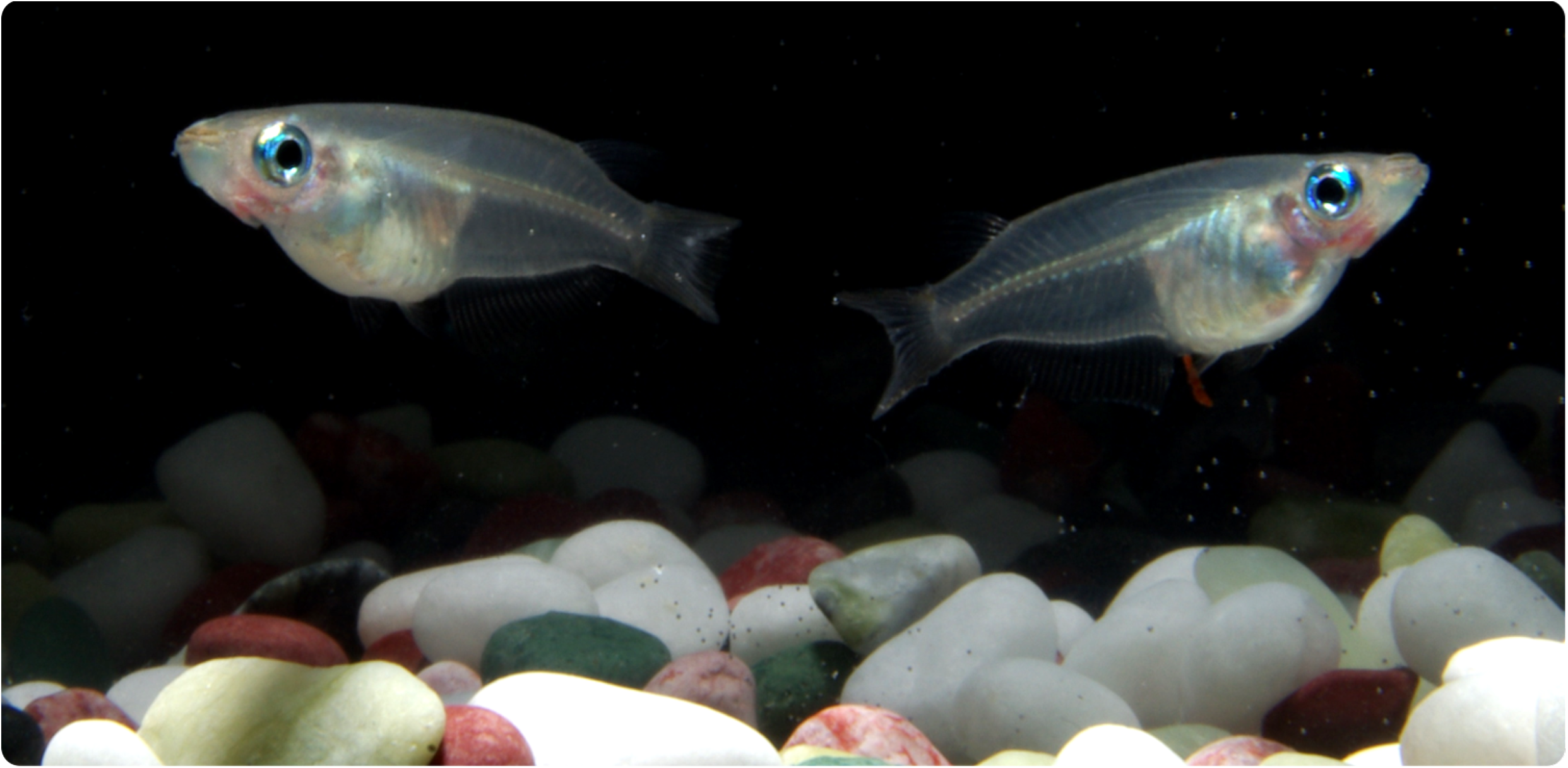
A couple of Java medakas, *Oryzias javanicus*. Picture from K. Naruse, NBRP Medaka stock centre (https://shigen.nig.ac.jp/medaka/top/top.jsp).

In this study, we sequenced and assembled the whole genome of *O. javanicus*, a model fish species for studying molecular mechanisms of seawater adaptation. In teleost fish, the major osmoregulatory organs i.e., gills, intestine and kidney, play different roles for maintaining body fluid homeostasis. Many genes encoding hormones, receptors, osmolytes, transporters, channels and cellular junction proteins are potentially involved in this osmotic regulation. In addition to osmoregulation, hatching enzyme activity dramatically fluctuates and adjusts at different salt conditions. At hatching stage, fish embryos secrete a specific cocktail of enzymes in order to dissolve the egg envelope, or chorion. In the medaka *O. latipes*, digestion of the chorion occurs through the cooperative action of two kinds of hatching enzymes, ***(i)*** the high choriolytic enzyme (HCE) and ***(ii)*** the low choriolytic enzyme (LCE) (19). The HCE displays a higher activity in fresh-than in brackish waters (20). Thus, availability of a high-quality reference genome in *O. javanicus* would facilitate further research for investigating the molecular basis of physiological differences, including the osmotic regulation and the hatching enzyme activity, among *Oryzias* species.

## SAMPLING AND SEQUENCING

### Animal samplings

The wild stock of *O. javanicus* used in this study was supplied by the National Bio-Resource Project (NBRP) medaka in Japan. This stock (strain ID: RS831) was originally collected at Penang, Malaysia, and maintained in aquaria under an artificial photoperiod of 14 hours light: 10 hours darkness at 27±2ºC. Genomic DNA was extracted from the whole body of a female (having ZW sex chromosome) using a conventional phenol/chloroform method, and was subjected to PacBio and 10X Genomics sequencings. For RNA-sequencing, total RNAs were extracted from nine female tissues (brain, bone, gill, heart, intestine, kidney, liver, muscle and ovary), and one male tissue (testis) using the RNeasy Mini Kit (Qiagen). For genetic mapping, we used a DNA panel consisting of 96 F1 progeny with their parents (originally described in a previous study (21)). Phenotypic sex was determined by secondary sex characteristics of adult fish, namely, the shapes of dorsal and anal fins. All animal experiments performed in this study complied with the guideline of National Institute for Basic Biology, and have been approved by the Institutional Animal Care and Use Committee of National Institute of Natural Science (16A050 and 17A048).

### Libraries construction and sequencing

#### PacBio genome sequencing

Library construction and sequencing were performed according to the manufacturer’s instructions (Shared protocol-20kb Template Preparation Using BluePippin Size Selection system (15kb size Cutoff)). When required, DNA was quantified using the Qubit dsDNA HS Assay Kit (Life Technologies). DNA purity was assessed by spectrophotometry using the nanodrop instrument (Thermofisher), and size distribution and absence of degradation were monitored using the Fragment analyzer (AATI) (8–11). Purification steps were performed using 0.45X AMPure PB beads (PacBio). 80μg of DNA was purified and then sheared at 40kb using the megaruptor system (diagenode). DNA and END damage repair step was further performed for 5 libraries using the SMRTBell template Prep Kit 1.0 (PacBio). Blunt hairpin adapters were then ligated to the libraries. Libraries were subsequently treated with an exonuclease cocktail in order to digest unligated DNA fragments. Finally, a size selection step using a 15kb cutoff was performed on the BluePippin Size Selection system (Sage Science) using 0.75% agarose cassettes, Marker S1 high Pass 15-20kb. Conditioned sequencing primer V2 was annealed to the size-selected SMRTbell. The annealed libraries were then bound to the P6-C4 polymerase using a ratio of polymerase to SMRTbell set at 10:1. After performing a magnetic bead-loading step (OCPW), SMRTbell libraries were sequenced on 48 SMRTcells (RSII instrument at 0.25nM with a 360-min movie resulting in a total of 61.8Gb of sequence data (1.28Gb/SMRTcell).

#### 10X Genomics genome sequencing

Chromium library was prepared according to 10X Genomics’ protocol using the Genome Reagent Kits v1. Sample quantity and quality controls were further validated on Qubit, Nanodrop and Femto. Optimal performance has been characterized on input gDNA with a mean length greater than 50 kb. The library was prepared using 3 μg of high molecular weight (HMW) gDNA (cut off at 50kb using BluePippin system). In details, for the microfluidic Genome Chip, a library of Genome Gel Beads was combined with HMW template gDNA in Master Mix and partitioning oil in order to create Gel Bead-In-EMulsions (GEMs) in the Chromium. Each Gel Bead was functionalized with millions of copies of a 10x™ Barcoded primer. Upon dissolution of the Genome Gel Bead in the GEM, primers containing (*i*) an Illumina R1 sequence (Read 1 sequencing primer), (*ii*) a 16 bp 10x Barcode, and (*iii*) a 6 bp random primer sequence were released. Read 1 sequence and the 10x™ Barcode were added to the molecules during the GEM incubation. P5 and P7 primers, Read 2, and Sample Index were added during library construction. 8 cycles of PCR were performed for amplifying the library. Library quality was assessed using a Fragment analyzer. Finally, the library was sequenced on an Illumina HiSeq3000 using a paired-end read length of 2×150 pb with the Illumina HiSeq3000 sequencing kits resulting in 101.6Gb of raw sequence data.

#### Transcriptome RNA-seq sequencing

RNA-seq libraries were prepared according to Illumina’s protocols using the Illumina TruSeq Stranded mRNA sample prep kit. Briefly, mRNAs were selected using poly-T beads, reverse-transcribed and fragmented. The resulting cDNAs were then subjected to adaptor ligation. 10 cycles of PCR were performed for amplifying the libraries. Quality of the libraries was assessed using a Fragment Analyser. Quantification was performed by qPCR using the Kapa Library Quantification Kit. RNA-seq libraries were sequenced on an Illumina HiSeq3000 using a paired-end read length of 2×150 pb with the Illumina HiSeq3000 sequencing kits resulting in 95Gb of sequence data (28.9M reads pairs/library).

#### RAD-library construction

RAD-seq library was built following the Baird et al. (22) protocol with minor modifications. Briefly, between 400 to 500 ng of gDNA per fish were digested with SbfI-HF enzyme (R3642S, NEB). Digested DNA was purified using AMPure PX magnetic beads (Beckman Coulters) and ligated to indexed P1 adapters (1 index per sample) using concentrated T4 DNA ligase (M0202T, NEB). After quantification (Qubit dsDNA HS assay kit, Thermofisher) all samples were pooled in equal amounts. The pool was then fragmented on a S220 sonicator (Covaris) and purified with Minelute column (Qiagen). Finally, the sonicated DNA was size selected (250 to 450 bps) on a Pippin HT (Sage science) using a 2 % agarose cassette, repaired using the End-It DNA-end repair kit (Tebu Bio) and adenylated at its 3’ ends using Klenow (exo-) (Tebu-Bio). P2 adapters were then ligated using concentrated T4 DNA ligase, and 50 ng of the ligation product were engaged in a 12 cycles PCR for amplification. After AMPure XP beads purification, the resulting library was checked on a Fragment Analyzer (Agilent) using the HS NGS kit (DNF-474-33) and quantified by qPCR using the KAPA Library Quantification Kit (Roche, ref. KK4824). Ultimately the whole library was denatured, diluted to 10 pM, clustered and sequenced using the rapid mode v2 SR100nt lane of a Hiseq2500 device (Illumina).

## Assembly results and quality assessment

### Genome Characteristics

To estimate size and other genome characteristics, 10X reads were processed with Jellyfish v1.1.11 (23) to produce 21-mer distribution. The k-mer histogram was uploaded to GenomeScope (24) with the max k-mer coverage parameter set to 10,000. Genome size was estimated around 908 Mbp, which is slightly higher than the 850 Mbp (0.87pg) estimated size reported on the Animal Genome Size Database (25). Furthermore, this analysis estimates that the *O. javanicus* genome contains 33% of repeat sequences (around 303 Mbp) and has a heterozygosity of 0.96% (Table 1).

**Table 1:** GenomeScope outputs on *O. javanicus* genome statistics.

| Property | min | max |
| --- | --- | --- |
| Heterozygosity | 0.960 % | 0.964 % |
| Genome Haploid Length | 908,146,324 bp | 908,641,143 bp |
| Genome Repeat Length | 303,610,795 bp | 303,776,222 bp |
| Genome Unique Length | 604,535,529 bp | 604,864,921 bp |
| Model Fit | 95.95 % | 99.72 % |
| Read Error Rate | 1.50 % | 1.50 % |

### Genome assembly with long PacBio reads and short 10X reads

PacBio reads were corrected and trimmed using Canu v1.5 (26). Contigs were then assembled using SMARTdenovo version of May 2017 (27). The draft assembly produced contains 729 contigs with a total genome size of 807.5 Mbp, an N50 of 3.9 Mbp and a L50 of 59 contigs (Figure 2). To improve the assembly base pair quality two polishing steps were run. First, BLASR aligned PacBio reads were processed with Quiver from the Pacific Biosciences SMRT link software v.4.0.0. Second, 10X reads were realigned to the genome using Long Ranger v2.1.1 and the alignment file was processed with Pilon v1.22 (28). Third, the same 10X reads were aligned to the genome with BWA-MEM v0.7.12-r1039 (29) and the alignment file was processed with ARCS v1.0.1 (30) to scaffold the genome. Both tools were run with default parameters. The final draft assembly contains 525 scaffolds with a total length of 809.7 Mbp, a N50 of 6.3 Mbp and a L50 of 37 scaffolds. This represents 89.1% of the k-mer estimated genome size. Given the high percentage of repeats in the *O. javanicus* genome (33%), it is possible that the PacBio assembly did not totally succeed in completing all repeated regions. The genome completeness was estimated using Benchmarking Universal Single-Copy Orthologs (BUSCO) v3.0 (31) based on 4,584 BUSCO orthologs derived from the Actinopterygii lineage leading to BUSCO scores of 4,327 (94.4%) complete BUSCOs, 176 (3.8%) fragmented BUSCOs and 81 (1.8%) missing BUSCOs.

**Figure 2:**
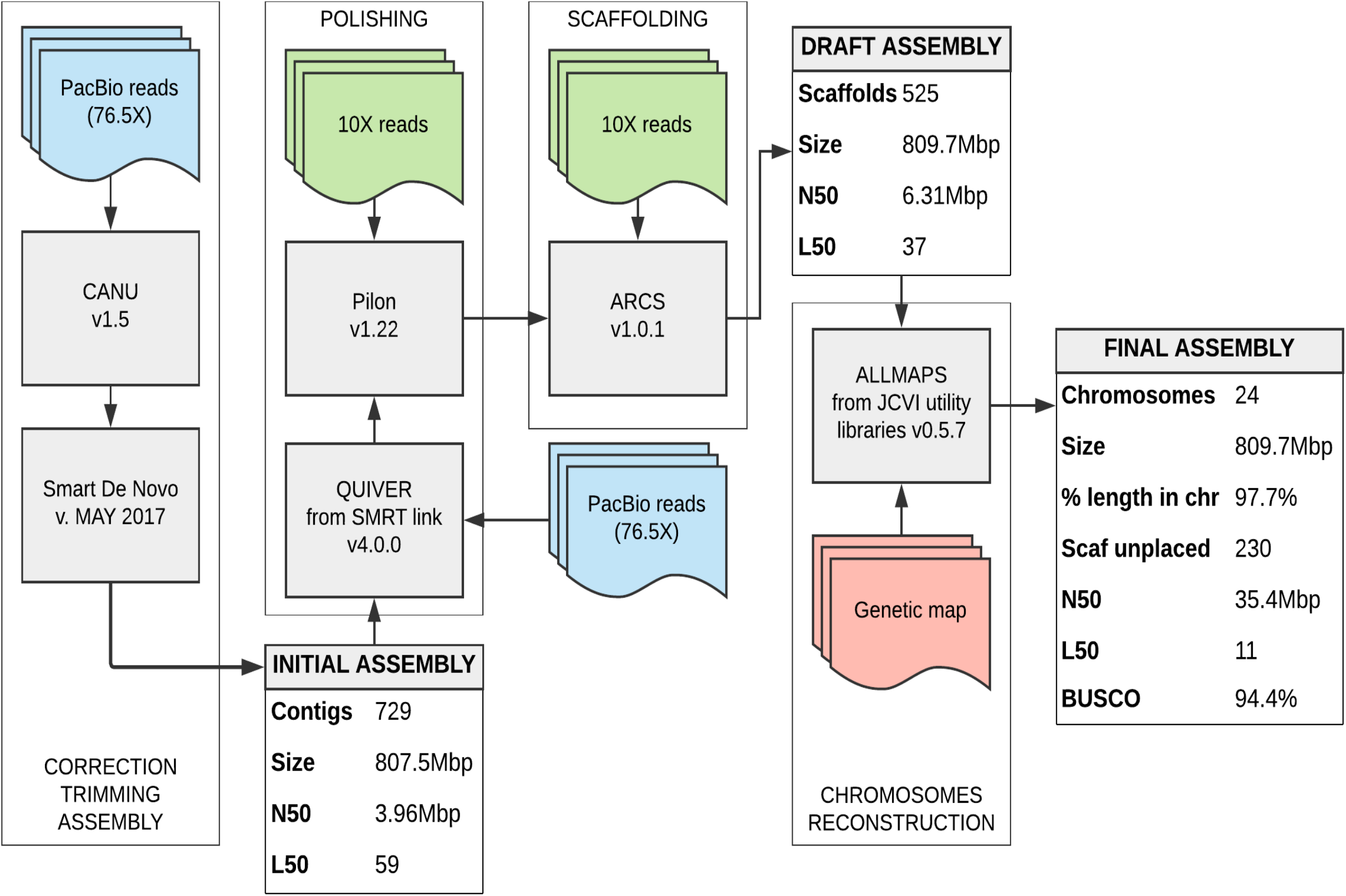
Oryzias javanicus assembly pipeline. Sequencing data are represented by coloured rectangles with waved bases. Tools used are in grey rectangles. Assembly metrics are in grey and white rectangles. This pipeline is divided in stages symbolized by the frame.

### Integration with the genetic map

RAD reads were trimmed by Trim Galore 0.4.3 (32) with Cutadapt 1.12 (33) and then mapped to the assembled scaffolds using BWA-MEM v0.7.17 (29). Uniquely mapped reads were extracted from the read alignments, and then called variant bases using uniquely mapped reads by samtools *mpileup* and bcftools *call* (34). Indels and variants with a low genotyping quality (GQ < 20), a low read depth (DP < 5), a low frequency of the minor allele (< 5%), more than four alleles in the family, no more than 5% individuals missing were removed by vcftools v0.1.15 (35). After quality filtering, 6,375 variant sites were kept for the following analysis. Linkage map was constructed using this genotype information using Lep-MAP3 (36). Briefly, the filtered vcf file was loaded and the markers removed with high segregation distortion (*Filtering2*: dataTolerance=0.001). Markers were then separated into 24 linkage groups with a LOD score threshold set at 9 and a fixed recombination fraction of 0.08 (*SeparateChromosomes2*: lodlimit=9 and theta=0.08). Two linkage groups were then excluded because of their small numbers of contained markers (less than 10). Classification of the markers was determined after maximum likelihood score indexing with 100 iterations (*OrderMarkers2*: numMergeIterations=100) in each linkage group. The final map had 5,738 markers dispatched amongst 24 linkage groups spanning a total genetic distance of 1,221 cM.

The linkage map exhibited discrepancies between genomic scaffolds and genetic markers. Among 525 genomics scaffolds, 32 were linked to more than one linkage group. To split chimeric scaffolds with a higher precision and to rebuild chromosomes with a higher fidelity, we used a cross-species synteny map between the Java medaka (*O. javanicus*) scaffolds and the medaka (*O. latipes*) chromosomes in order to combine marker locations from genetic and synteny maps. To build the synteny map, medaka cDNAs were aligned to the Java medaka scaffolds using BLAT v36 (37), and a list of pairwise correspondence of gene positions on Java medaka scaffolds and medaka chromosomes was established. 13,796 markers were added to the 5,738 markers of the genetic map. Java medaka chromosomes were then reconstructed using ALLMAPS from the JCVI utility libraries v0.5.7 (38). This package was used to combine genetic and synteny maps, to split chimeric scaffolds, to anchor, order and orient genomic scaffolds. The resulting chromosomal assembly consists of 321 scaffolds anchored on 24 chromosomes (97.7% of the total bases) and 231 unplaced scaffolds

### Transcriptome assembly

The read quality of the RNA-seq libraries was evaluated using FastQC (39). *De novo* and reference-based transcriptome assemblies were produced. Reads were cleaned, filtered and *de novo* assembled using the DRAP pipeline v1.91 (40) with the Oases assembler (41). Assembled contigs were filtered in order to keep only those with at least one fragment per kilobase of transcript per million reads (FPKM). In the reference-based approach, all clean reads were mapped to the chromosomal assembly using STAR v2.5.1b (42) with outWigType and outWigStrand options to output signal wiggle files. Cufflinks v2.2.1 (43) was used to assemble the transcriptome.

### Annotation results

The first annotation step was identifying repetitive DNA content using RepeatMasker v4.0.7 (44), Dust (45) and TRF v4.09 (46). A species-specific *de novo* repeat library was built with RepeatModeler v1.0.11 (47). Repeated regions were located using RepeatMasker with the *de novo* and the Zebrafish (*Danio rerio*) libraries. Bedtools v2.26.0 (48) was used to merge repeated regions identified with the three tools and to soft mask the genome. Repeats were estimated to account for 43.16% (349 Mbp) of our chromosomal assembly. The MAKER3 genome annotation pipeline v3.01.02-beta (49) combined annotations and evidences from three approaches: similarity with known fish proteins, assembled transcripts and *de novo* gene predictions. Protein sequences from 11 other fish species found in Ensembl were aligned to the masked genome using Exonerate v2.4 (50). Previously assembled transcripts were used as RNA-seq evidence. A *de novo* gene model was built using Braker v2.0.4 (51) with wiggle files provided by STAR as hints file for training GeneMark and Augustus. The best supported transcript for each gene was chosen using the quality metric Annotation Edit Distance (AED) (52). The genome annotation gene completeness was assessed by BUSCO using the Actinopterygii group (Table 2). Finally, the predicted genes were subjected to similarity searches against the NCBI NR database using Diamond v0.9.22 (53). The top hit with a coverage over 70% and identity over 80% was retained.

**Table 2:** Java medaka assembly and annotation statistics.

| Gene annotation |  |
| --- | --- |
| Number of genes | 21,454 |
| Number of transcripts | 21,454 |
| Transcriptome size | 57,146,583 bp |
| Mean transcript length | 2,663 bp |
| Longest transcript | 42,733 bp |
| Number of genes with significant hit against NCBI NR | 17,412 (81.2%) |
| Gene completeness |  |
| Complete BUSCOs | 4,289 (93.6%) |
| Fragmented BUSCOs | 187 (4.1%) |
| Missing BUSCOs | 108 (2.3%) |

### Mitochondrial genome and annotation

The previously sequenced *Oryzias javanicus* mitochondrial genome (NC_012981) (54) was aligned to the chromosomal assembly using Blat. All hits were supported by a single scaffold. This scaffold was removed from the assembly, circularised and annotated using MITOS (55). This new *Oryzias javanicus* mitochondrial genome is 16,789 bp long and encodes 13 genes, 2 rRNAs and 19 tRNAs.

### Phylogenetic relationship

To precisely determine the phylogenetic position of *O. javanicus* within the genus *Oryzias*, we estimated the phylogenetic relationship using published whole genome datasets as references. Reference assemblies and annotations of *O. latipes* (Hd-rR: ASM223467v1), *O. sakaizumii* (HNI-II: ASM223471v1), *Oryzias* sp. (HSOK: ASM223469v1), *O. melastigma* (Om_v0.7.RACA), and southern platyfish *Xiphophorus maculatus* (X_maculatus-5.0-male) were obtained from Ensembl Release 94 (http://www.ensembl.org/). Among the six genomes, orthologous groups were classified and 10,852 single-copy orthologous genes were identified using OrthoFinder 2.2.6 (56). For every single gene, codon alignment based on translated peptide sequences was generated by PAL2NAL (57) and then trimmed by trimAl with ‘-autometed1’ option (58). All multi-sample fasta files were concatenated into a single file using AMAS *concat* by setting each gene as a separate partition (59). A maximum likelihood tree was then inferred using IQ-TREE v1.6.6 (60) with the GTR+G substitution model for each codon, followed by an ultrafast bootstrap analysis of 1,000 replicates (61). This tree (Figure 3) indicates that *O. javanicus* forms a monophyletic group with *O. melastigma* but not with the *O. latipes* species complex (Hd-rR, HNI-II, and HSOK), being consistent with previous trees inferred from two mitochondrial genes and a nuclear gene (14).

**Figure 3:**
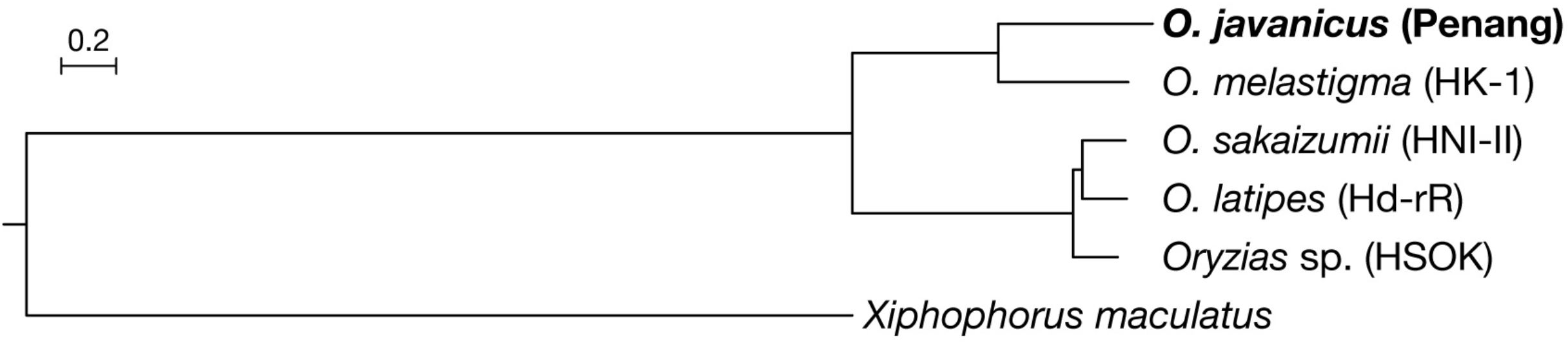
Phylogenetic position of *O. javanicus*. Maximum likelihood tree was inferred from the concatenated codon-alignment of 10,852 single-copy genes among 5 reference assemblies of *Oryzias* species with Southern platyfish (*Xiphophorus maculatus*) as outgroup. All nodes were supported by 100% bootstrap values.

### Adaptation to salinity and hatching enzymes

To gain insight into gene family evolution associated with osmoregulation, we used HMMER version 3.1b2 (62) to identify Pfam domain (Pfam 32, El-Gebali et al., 2019) containing proteins in the *O. javanicus* genome. We used protein sequences based on our gene model of *O. javanicus* combined with Ensembl genes of the *O. latipes* species complex (Hd-rR, HNI-II and HSOK) and *O. melastigma* (a synonym of *O. dancena*) for the Pfam search, and focused on 147 domains found in 224 proteins whose functions were related to osmoregulation (Additional Tables 1 and 2). Similar numbers of proteins were observed among species for each domain, suggesting that the osmoregulation gene repertoires are relatively conserved in *Oryzias* species. However, further detailed comparisons are required because gene annotation methods are different among data.

We then also focused on specific genes encoding hatching enzymes. In the genome of *O. latipes*, five copies of HCE genes-including one pseudogene-are clustered tandemly with the same transcriptional direction on chromosome 3 (chr. 3), while only one single copy of the LCE gene is located on chromosome 24 (chr. 24) (63). In *O. javanicus* 5 copies of the HCE (OjHCE) gene are located on chromosome 3 and one LCE (OjLCE) gene was found on chromosome 24. The amino acid sequence similarities in the mature enzyme region of the 5 OjHCE genes are between 89-99%. Only in comparison to *O. latipes*, within the five *O. javanicus* HCE genes, the fourth one (OjHCE4) displays an opposite orientation compared to the others (Figure 4A) suggesting a rearrangement within the HCE gene cluster that has likely been occurring during the evolution of *Oryzias* lineage.

**Figure 4.**
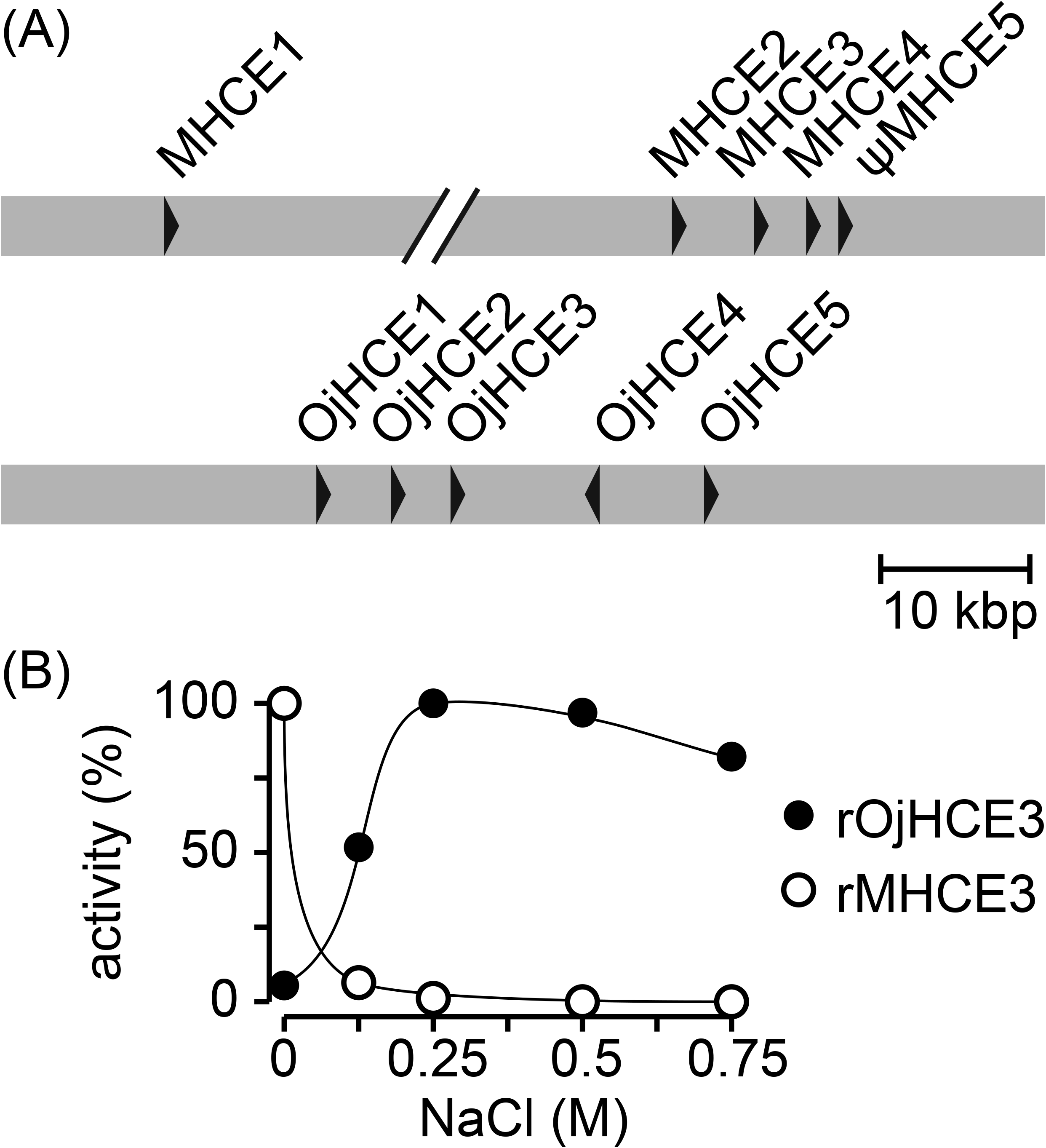
Hatching enzyme of Oryzias javanicus. (A) HCE gene cluster of *O. latipes* (MHCE1-5) and *O. javanicus* (OjHCE1-5). Arrowheads indicate direction of transcription. (B) Salt dependency of *O. javanicus* HCE (black circle) and *O. latipes* (white circle). Activities are shown as % of relative activity with respect to highest activity, which is considered as 100% in each species.

While LCE’s activity remains constant over various salinities, HCEs have been reported to show salt-dependent activity (49). In contrast to other *Oryzias* species, *O. javanicus*, being a euryhaline species, specifically adapted its physiology to higher water salinities. In order to test whether such adaptive evolution would translate at the level of HCE activity, recombinant OjHCE3 (rOjHCE3) was generated in an *E. coli* expression system, refolded, and its activity regarding to the digestion of the egg-envelope determined at various salt concentrations based on the method described in Kawaguchi et al. (49). Although rOjHCE3 showed virtually no activity at 0 M NaCl, an increased activity was apparent at elevated salt concentrations. Furtheron rOjHCE3 activity was recorded to be highest at 0.25 M NaCl, while still maintaining high activity up to 0.75 M NaCl (Figure 4B). In contrast, it has been reported that *O. latipes* HCEs show highest activity at 0 M NaCl, and drastically decrease when salt concentrations increase ((20), Figure 4B). These results suggest that salt preference of HCE enzymes is a species-specific adaptation to different salt environments at hatching.

## CONCLUSION

The Java medaka, *Oryzias javanicus*, is one of the two species of medaka perfectly adapted for living in brackish/sea-waters. Being an important component of the mangrove ecosystem, *O. javanicus* is also used as a valuable marine test-fish for ecotoxicology studies. Here, we sequenced and assembled the whole genome of *O. javanicus*. Complementary sequencing approaches and data integration with a genetic map allowed the final assembly of the 908 Mbp of the *O. javanicus* genome. The final draft assembly contains 525 scaffolds with a total length of 809.7 Mbp, a N50 of 6.3 Mbp and a L50 of 37 scaffolds. Providing here a high-quality draft genome assembly of the euryhaline Javafish medaka, we anticipate this resource will be catalytic for a wide range of comparative genomic, phylogenetic and functional studies within the genus *Oryzias* and beyond.

## Availability of supporting data

All genome and transcriptome datasets are available at the GigaDB repository [XX]. The genome assembly has also been deposited at GenBank under whole genome shotgun sequencing project accession number RWID00000000.1. Illumina genome and transcriptomes and PacBio genome raw reads are also available in the Sequence Read Archive (SRA), under BioProject reference PRJNA505405.

## Author contributions

-Designed the project: YT, AH, YG, KN

-Collected the samples and prepared the quality control: YT

-Sequencing data production: CR, OB, CD, CB, LJ

-Data analysis: MZ, CC, CK, MK, SY, SA, KI, CS

-Wrote the manuscript: YT, AH, MK, MS, SY, SA

-Supervision, project administration and funding acquisition: YT, AH, YG, MK, KN

All the authors read and approved the final manuscript.

## Acknowledgements

This work was supported by a “Projet Incitatif PHASE department 2015” grant (Grant ID ACI_PHASE, Institut National de la Recherche Agronomique) to AH, a NIBB Collaborative Research Initiative to KN, NIBB individual Collaboration Research Project to MK, and Grants-in-Aid for Young Scientists to YT (Grant ID 16K18590) and MK (Grant ID 16K18593). The GeT and MGX core facilities were supported by France Génomique National infrastructure, funded as part of “Investissement d’avenir” program managed by Agence Nationale pour la Recherche (contract ANR-10-INBS-09). The GeT core facility was also supported by the GET-PACBIO program (« Programme operationnel FEDER-FSE MIDI-PYRENEES ET GARONNE 2014-2020 »).

